# Differentiation of Urine-Derived Induced Pluripotent Stem Cells to Neuron, Astrocyte and Microvascular Endothelial Cells from a Diabetic Patient

**DOI:** 10.1101/667485

**Authors:** Wan Liu, Ping Zhang, Jing Tan, Yongzhong Lin

**Affiliations:** Department of Neurology, the Second Hospital of Dalian Medical University, No. 467 Zhongshan Road, Shahekou District, Dalian City 116027, Liaoning Province, China; Department of Endocrinology, the Second Hospital of Dalian Medical University, No. 467 Zhongshan Road, Shahekou District, Dalian City 116027, Liaoning Province, China

**Keywords:** Human induced pluripotent stem cells, Urinary epithelial cells, Diabetes mellitus, Cognitive impairment, Personalized disease model, Precision medicine

## Abstract

**Background:** Complications of central nervous system (CNS) in type 2 diabetes mellitus (T2DM) often lead to cognitive impairment and seriously affect the quality of life. However, there is no individualized disease model. Urinary epithelial cells (UECs) can be an ideal source for generating human induced pluripotent stem cells (hiPSCs) and progenitors, as they are easily accessible, non-invasive and universally available. Therefore, we intended to differentiate urine-derived hiPSCs into neuron (N), astrocyte (A) and microvascular endothelial cells (E) from a T2DM patient for future study its pathogenesis and precision medical treatment.

**Methods and Results:** hiPSCs was successfully induced from UECs using integration free Sendai virus technology in a totally noninvasive manner. It had a normal karyotype (46, XY) and were proved to be pluripotent by immunofluorescence staining, alkaline phosphatase staining, karyotyping, teratoma experiments and methylated analysis. N, A and E were successfully induced and displayed typical morphological characteristics.

**Conclusions:** This study indicates that N, A, E can be generated from urine-derived hiPSCs. Then we intend to create a new disease model in vitro to simulate the cerebral microenvironment of DM which will provide new methods for further investigate the disease-specific mechanisms.

## Introduction

The injures of diabetes mellitus (DM) in the central nervous system (CNS) are mainly characterized by diabetes-induced cognitive impairment, neurophysiological and structural changes of the brain, such as vascular dementia (VD), cognitive decline, depression, epilepsy. In the past 20 years, the cognitive ability of middle-aged diabetics has decreased by 19% compared with those without diabetes(Mayeda *et al*., 2015). Especially, substantial evidence is accumulating that type 2 diabetes mellitus (T2DM) are related to cognitive impairment, and T2DM is an established risk factor for developing dementia and Alzheimer’s disease (AD)(Biessels *et al*., 2006a, Kopf and Frolich, 2009, Albai *et al*., 2019). In addition, the risk of AD or VD in patients with T2DM is 1.5-2 times higher than that in normal people(Biessels *et al*., 2006b). However, further studies related to molecular mechanisms of brain injury with DM have not been well-elucidated. In the preceding work of our research group(Wang *et al*., 2014, Lin *et al*., 2013), we found that the occurrence, performance, treatment and prognosis of T2DM brain injuries, such as cognitive impairment and hemichorea, had complicated and personalized relationship with the neuron (N), astrocyte (A) and microvascular endothelial cell (E) and other target cells, the cognitive impairment of diabetes mellitus was mainly related to the decrease of neurons in frontal lobe and hippocampus, as well as the proliferation of astrocytes and the destruction of blood-brain barrier. However, the factors and molecular signaling pathways that cause N, A and E cell injury/activation are still unclear. At present, most studies on diabetic brain injury used animal disease models, which will take a long time and have a low success rate, and there are some differences between biomedical animal models and human disease. There is no individualized model for further study of its pathogenesis.

Stem cells represent a great versatile cell source, as they are able to undergo a very high number of divisions because of their self-renewal property and furthermore differentiate into almost all adult cell types thanks to their pluripotency characteristic(Bordoni *et al*., 2018). Induced pluripotent stem cells (iPSCs) own similar properties with embryonic stem cells (ESCs) in terms of self-renewal, pluripotency, gene expression, proliferation, morphology, and telomerase activity(Takahashi *et al*., 2007, Takahashi and Yamanaka, 2006). In the past decades, the human induced pluripotent stem cells (hiPSCs) have found revolutionary application on vitro models of various neurological diseases, such as Alzheimer’s disease (AD), Parkinson disease (PD) and amyotrophic lateral sclerosis (ALS)(Iovino *et al*., 2015, Dimos *et al*., 2008, Soldner *et al*., 2009). In addition, hiPSCs can circumvent the ethical concerns and immune rejection which obstructed the application of human ESCs, and eliminate the shortcomings of species differences and individualized differences in animal models. Its unique potential of self-renewal makes it an exciting candidate for cell replacement therapy (CRT) for various diseases, such as neurodegenerative diseases and cancers, and offers unlimited possibilities for understanding early disease development and establishing in vitro disease models(Dakhore *et al*., 2018). Urinary epithelial cells (UECs) is an ideal source of iPSCs as it can be collected by a safe and negligibly invasive way and retain up to 0.1-4% of reprogramming ability that is much more than blood cells and fibroblasts(Raab *et al*., 2014). Furthermore, repeated freeze-thaw cycle will not exert an influence on its reprogramming ability(Raab et al., 2014). Sauer et al(Sauer *et al*., 2016) and Wang et al(Wang *et al*., 2016) reported human UECs could be efficiently reprogrammed into authentic iPSCs and further differentiated into targeted cell types, as well as opened the door for great potentials to promote regenerative therapies into the clinic.

Thus, we hypothesized that reprogramming of Urine-derived iPSCs into N, A and E may be a new noninvasive strategy for the study molecular mechanisms of disease pathogenesis, test potential drug toxicology reaction, and develop personalized therapeutic strategies for diabetic brain injury.

## Materials and methods

### Establishment of hiPSC line

#### Urinary cell isolation and expansion

After obtaining written informed consents from the donors, the urine samples were obtained from a 54-year old male patient with T2DM in the Second Hospital of Dalian Medical University. 300-500 ml of fresh midstream urine was collected in sterile containers, then centrifuged at 400rpm for 10 mins, and washed with 10 ml phosphate-buffered saline (PBS). After repeating twice of the above steps, the cell pellet was resuspended in 0.5 ml of Urineasy full-medium I (cellapybio, China) and maintained in Matrix coated plates. During the next 3-7 days, half of the medium was replaced with fresh medium every two days as follows: suctioned 2 ml medium, centrifuged at 200×g for 5 mins, the supernatant was discarded leaving 200 ul, added 2 ml Urineasy full-medium II, and then putted it back to the plates. Replaced the medium every two days with 3 ml Urineasy full-medium II after the cell successively attached. After reaching about 90% confluency, cells were passaged in the appropriate ratio (Fig.1).

**Fig. 1.**
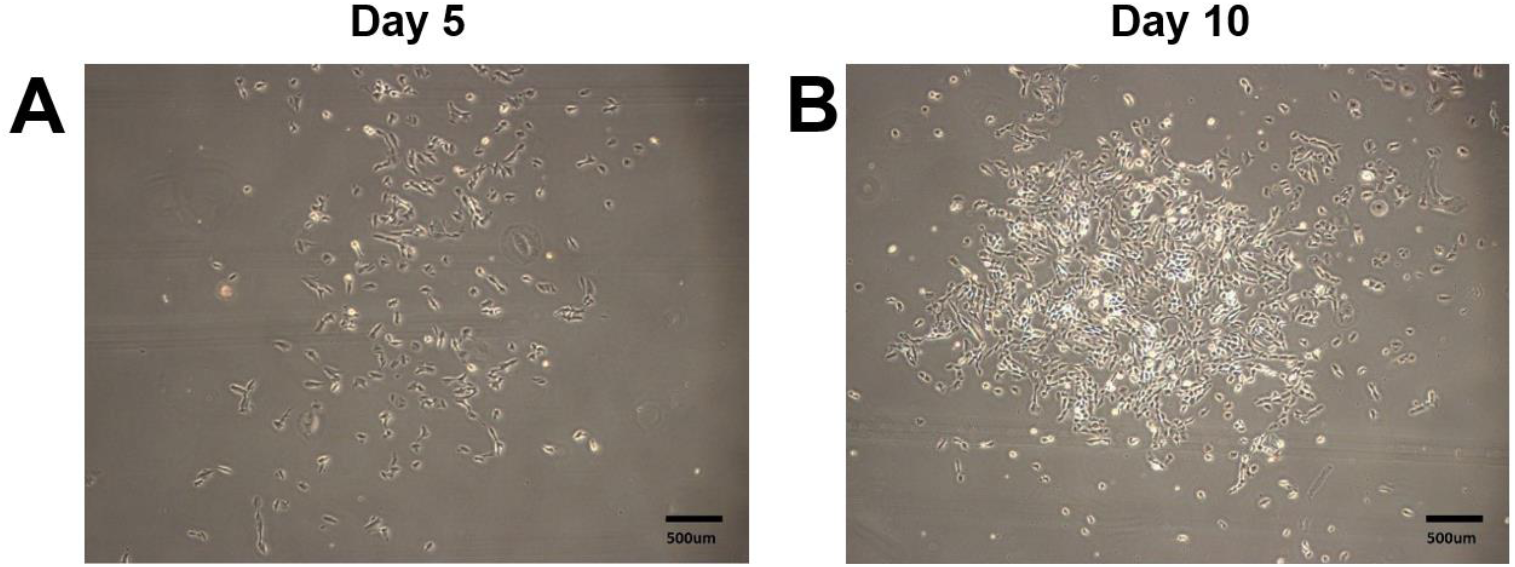
Urinary cell isolation and expansion. (A) Urinary cells attached to the plate in about 3 to 7 days. (B) Urinary cells reached about 90% confluency. Scale bars:500 µm.

#### Reprogramming of human urine renal epithelial cells

To generate hiPSC line, the reprogrammed urine cells were cultured in Urineasy full-medium II until reaching confluence of about 80%. According to the manufacturer’s instructions, these cells were reprogrammed into pluripotent stem cells by using a non-integrated iPS 2.0 Sendai Reprogramming Kit —CytoTune™. The medium was changed every two days. The transfected cells were inoculated in a culture dish coated with Matrigel containing urine maintenance medium on day 7. The plates were observed for the emergence of characteristic cell colonies every day. Around 3–4 weeks after transfection, picked undifferentiated colonies and transferred them onto fresh Matrigel-coated plates for further expansion and analysis (Fig.2).

**Fig. 2.**
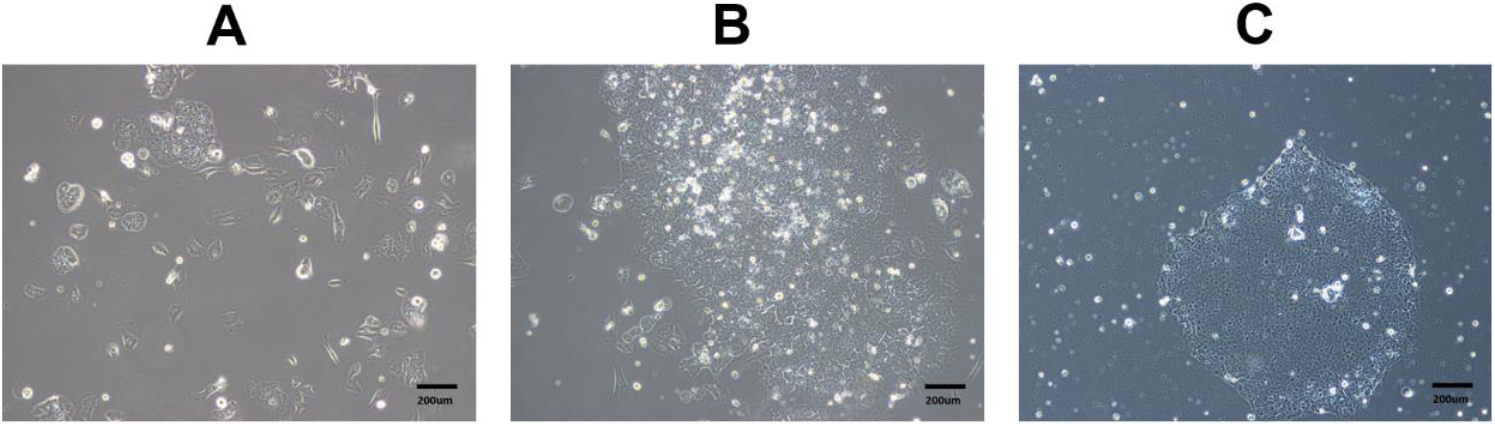
Reprogramming of human urine renal epithelial cells. (A) Characteristic cell colonies appeared. (B) Undifferentiated colonies after transfection at around 3–4 weeks after. (C) Cell expanded to the fifth generation. Scale bars:200 µm.

### Identification of pluripotency of hiPSCs cell line

#### Immunofluorescence staining

Immunofluorescence staining was used to analyze the expression of specific multipotent markers. The methods were slightly similar to that of Uhm KO (Uhm *et al*., 2017) and Zhang et al(Zhang *et al*., 2018). Cells were fixed in 4% paraformaldehyde for 30 mins, permeabilized with 0.3% Triton X-100 (Sigma, USA) for 20 mins, blocked in 1% bovine serum albumin for 40 mins and incubated with primary antibodies (Table 1) for TRA-1-60, TRA-1-81 and SSEA-4. Cell nuclei were labeled with 4′,6-diamidino-2-phenylindole (DAPI, Vector). Images were acquired using a fluorescence microscope (Carl ZeiSS, Germany) (Fig.3).

**Table 1.**
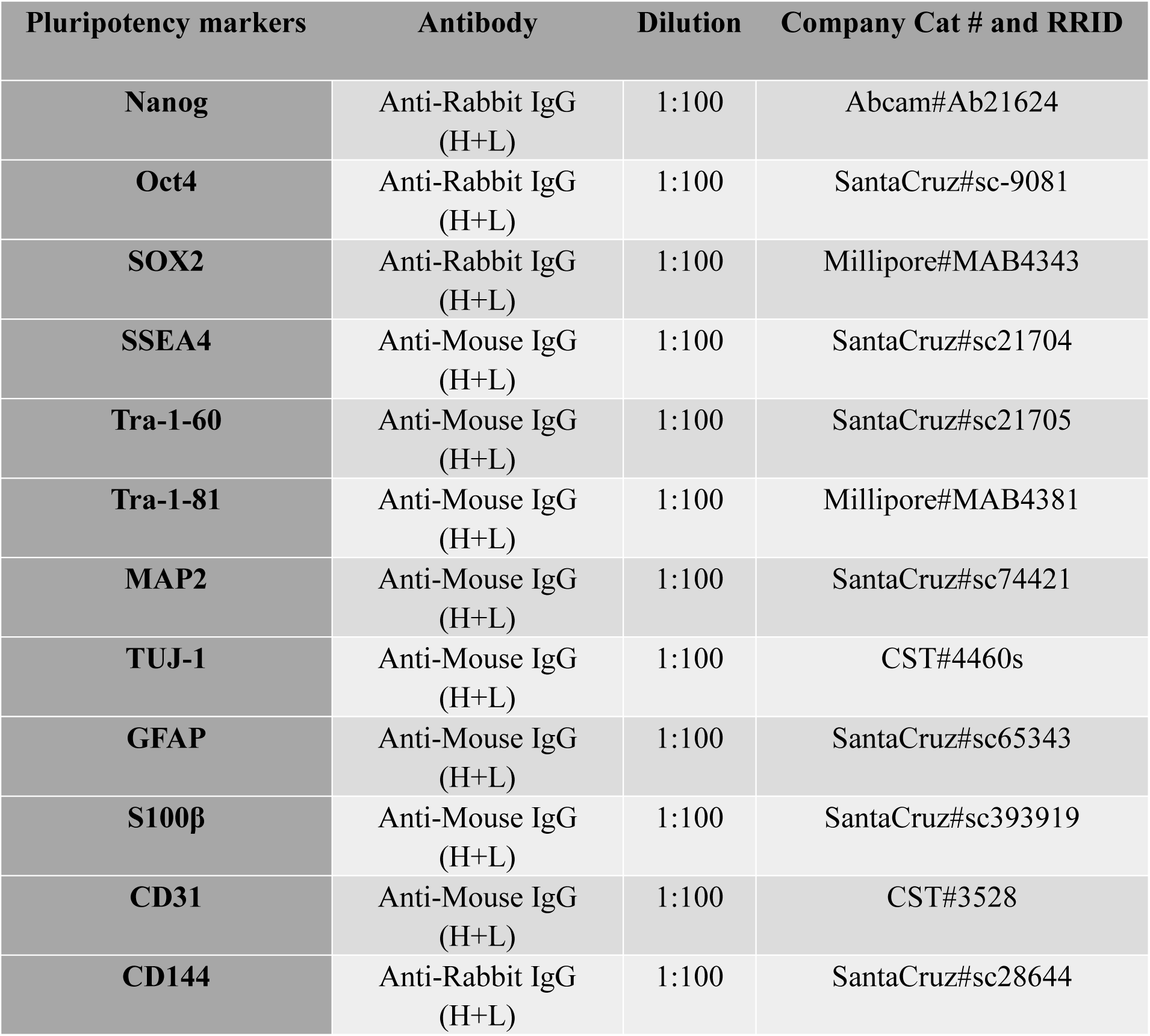
Antibodies used for immunocytochemistry.

| Pluripotency markers | Antibody | Dilution | Company Cat # and RRID |
| --- | --- | --- | --- |
| **Nanog** | Anti-Rabbit IgG (H+L) | 1:100 | Abcam#Ab21624 |
| **Oct4** | Anti-Rabbit IgG (H+L) | 1:100 | SantaCruz#sc-9081 |
| **SOX2** | Anti-Rabbit IgG (H+L) | 1:100 | Millipore#MAB4343 |
| **SSEA4** | Anti-Mouse IgG (H+L) | 1:100 | SantaCruz#sc21704 |
| **Tra-1-60** | Anti-Mouse IgG (H+L) | 1:100 | SantaCruz#sc21705 |
| **Tra-1-81** | Anti-Mouse IgG (H+L) | 1:100 | Millipore#MAB4381 |
| **MAP2** | Anti-Mouse IgG (H+L) | 1:100 | SantaCruz#sc74421 |
| **TUJ-1** | Anti-Mouse IgG (H+L) | 1:100 | CST#4460s |
| **GFAP** | Anti-Mouse IgG (H+L) | 1:100 | SantaCruz#sc65343 |
| **S100β** | Anti-Mouse IgG (H+L) | 1:100 | SantaCruz#sc393919 |
| **CD31** | Anti-Mouse IgG (H+L) | 1:100 | CST#3528 |
| **CD144** | Anti-Rabbit IgG (H+L) | 1:100 | SantaCruz#sc28644 |

#### Alkaline phosphatase staining

BCIP/NBT Alkaline Phosphatase Color Development Kit was used to analyze the expression of specific pluripotency. Cells were incubated with antibodies labeled with alkaline phosphatase, washed 3-5 times with PBS for 3-5 mins each time, added BICP/NBT staining solution, incubated in dark for 5-30 mins, and then washed twice with distilled water. AP stained (red/purple) pluripotent colonies and the differentiated (clear) colonies were observed by a light microscope with 10x objective. (Fig.4)

**Fig. 3.**
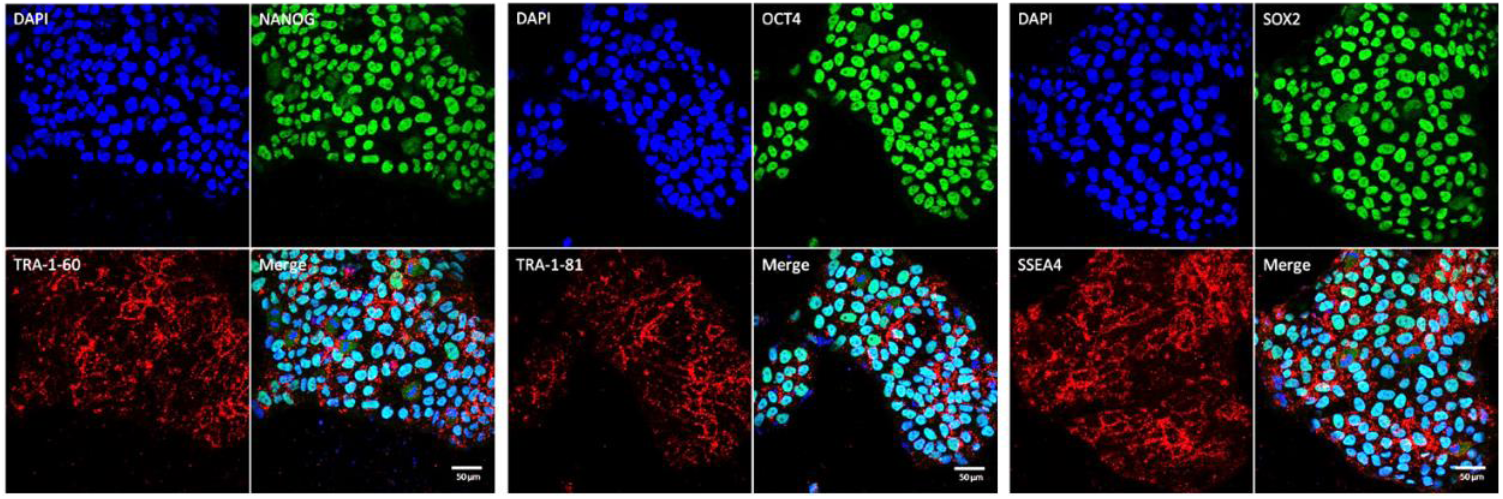
Immunofluorescence staining of hiPSCs. The expression of human embryonic stem cell (hESC)-specific transcription factors NANOG, OCT4 and SOX2, as well as cell surface markers SSEA4, TRA-1-60 and TRA-1-81 were observed. hiPSCs lines showed positive immunofluorescence labelling for above antigens. Scale bars:50 µm.

**Fig. 4.**
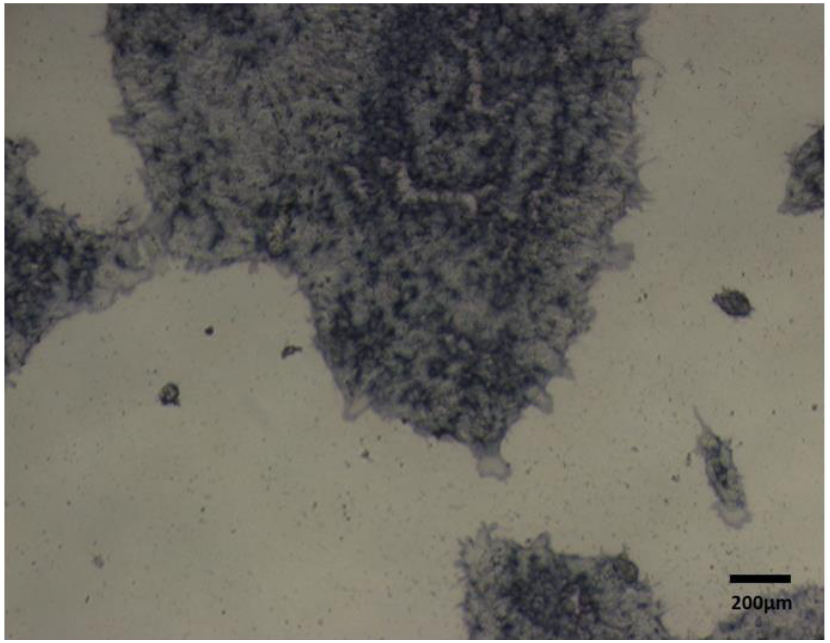
Alkaline phosphatase staining of hiPSCs. The pluripotency of hiPSC colonies displayed positive reaction for AP staining. Scale bars:200 µm.

#### Detection of reprogramming vector

After 10 passages, reprogrammed hiPSC lines were tested for SeV residues. The RT-PCR was performed as standard protocol. PCR was performed using primers (Table 2) and instructions as recommended by the manufacturer. As positive control RNA was used from the reprogramming leftovers. Negative control RNA was obtained from the Embryonic stem cell line H9 (Fig.5).

**Table 2.**
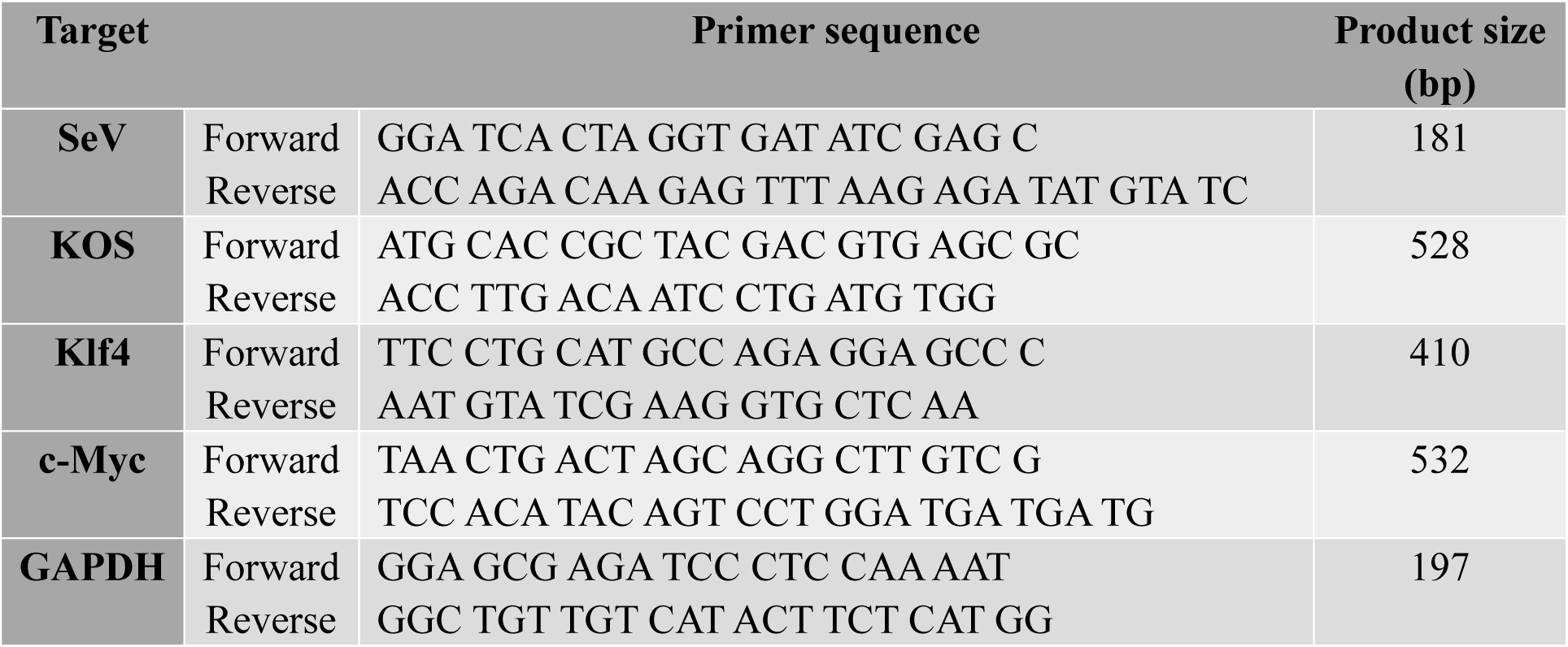
Primer sequences for Sendai viral test.

**Fig. 5.**
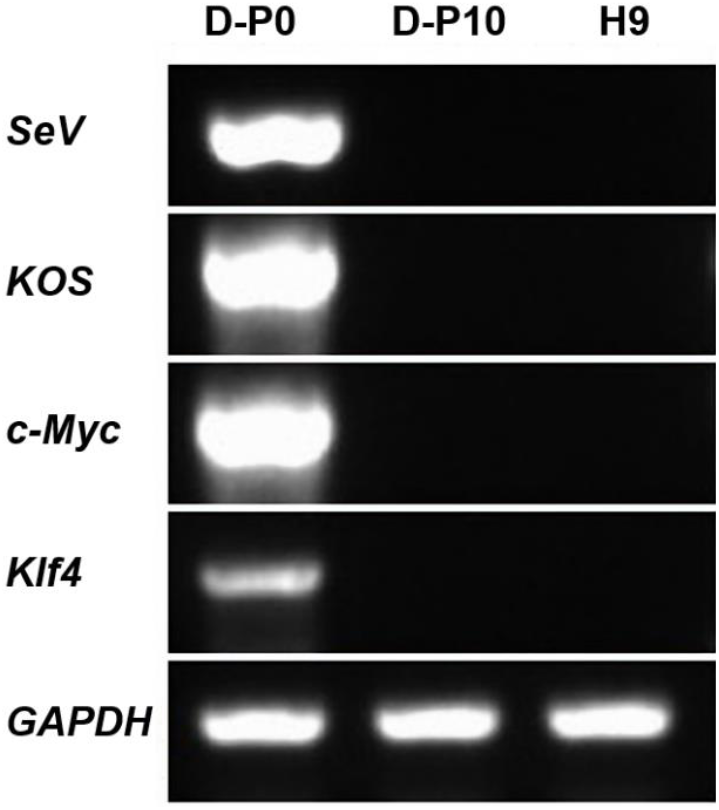
The absence of Sendai-virus from the hiPSCs lines were defected by RT-PCR, H9 ESCs was used as negative control. Scale bar: 10 µm.

#### Karyotyping

GTG-band method was used to analyze the karyotype of hiPSCs by standard cytogenetic procedures. Then cells were treated with colcemid (50 ng/ml) for 6-8 h, incubated in KCL hypotonic solution at 37 °C for 20–40 mins, fixed with fresh methanol/glacial acetic acid (3:1). The karyotype was detected and analysed using the VideoTesT-Karyo 3.1 system (Fig.6).

**Fig. 6.**
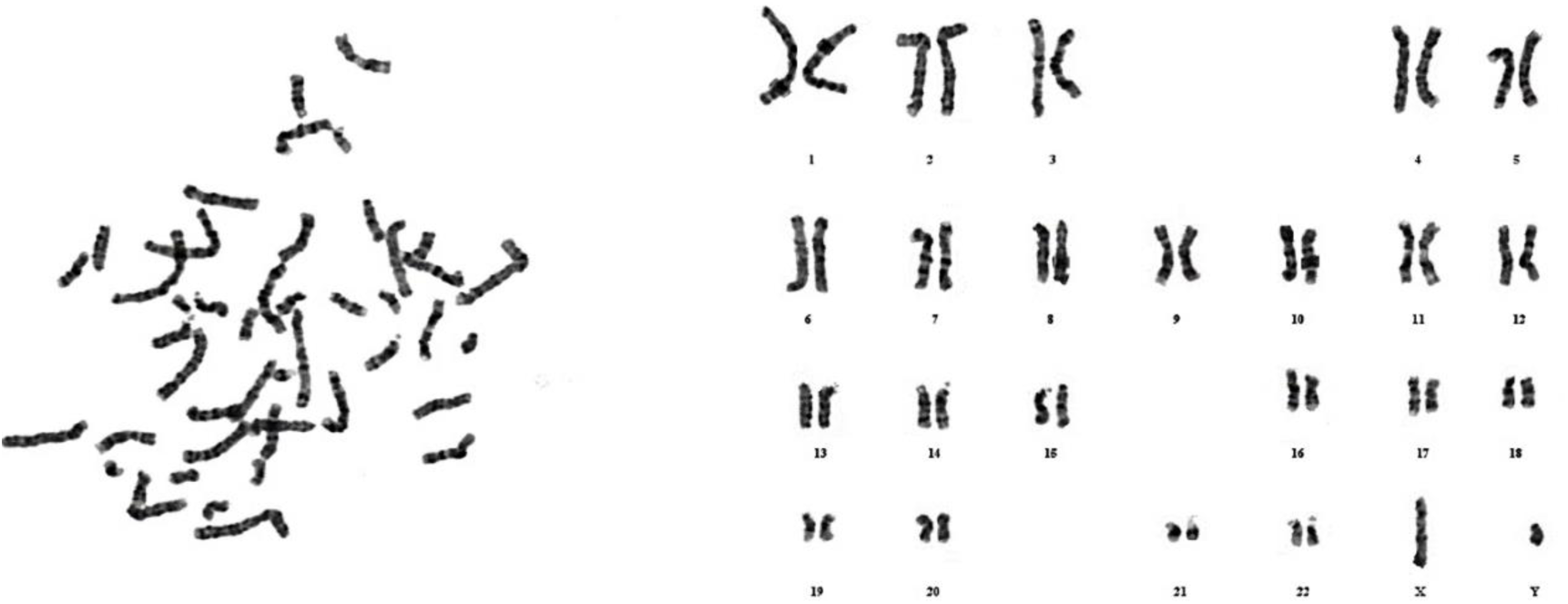
Karyotype analysis of hiPSCs lines showed 46 normal male chromosomes (XY).

#### Teratoma test

To test the capacity of the reprogrammed hiPSC line to spontaneously differentiate into cells of all three germ layers, the formation and differentiation of embryoid body (EB) were analyzed. We injected 1 × 10^7^ reprogrammed hiPSC cells into 4-week-old CB-17 SCID male mice to generate teratomas. The tumors were excised after 2-3 months, fixed in 4% PFA for 24 hours and embedded in paraffin. The embedded paraffin block was cut into 10um, oven at 60 °C for 1 hour and baking at 75 °C for 2-3 hours. Then hematoxylin/eosin staining was used to observe the formation of the three germ layers (Fig.7).

**Fig. 7.**
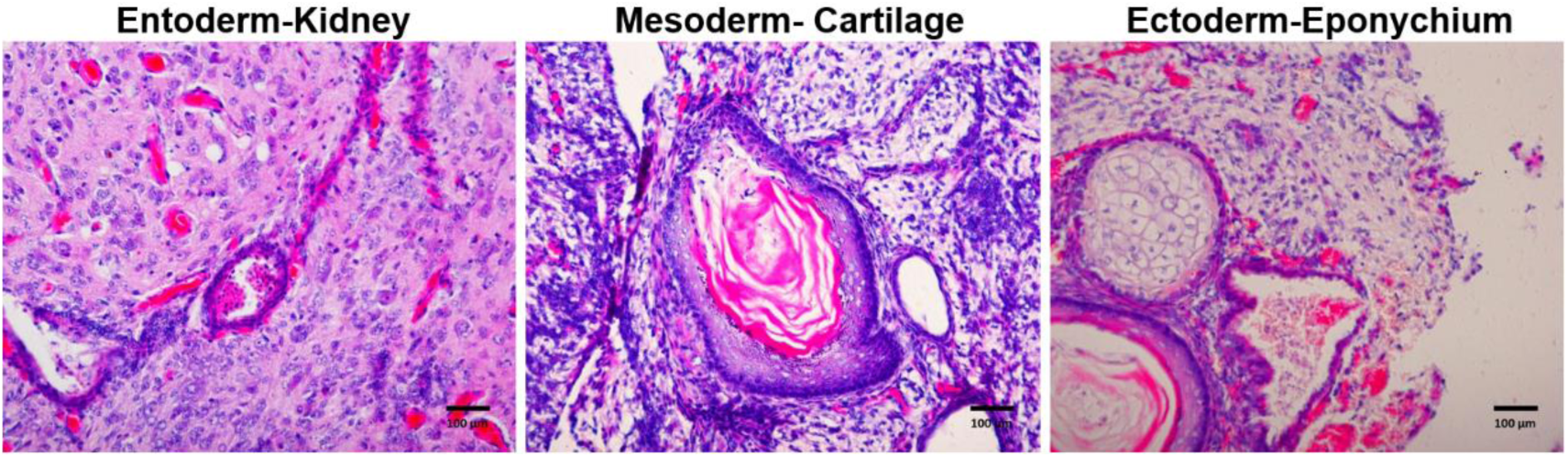
Hematoxylin/eosin staining of teratoma tissue sections produced by hiPSC cloning. All three germ layers could be detected in vivo. Scale bar: 100 µm.

#### Methylation analysis

We used Bisulfite sequencing PCR(BSP) for methylation analysis. Genomic DNA was treated with bisulfite, and then primers were designed on the periphery of CpG island (excluding CpG loci) for PCR. According to gene sequence information, the primers (Table 3) were designed and synthesized by primer5 software. Finally, the PCR products were sequenced. The CPG methylation of OCT 4 and NAG promoters were determined by Real-time quantitative PCR (qRT-PCR) on the GeneAmp 9600 PCR assay system (ABI Bio-Rad, Hercules, USA). PCR cycle was used as follows: 96°C for 2 mins, followed by 30 cycles of 95°C for 15 s, 50°C for 20 s, and 60°C for 4 mins (Fig.8).

**Table 3.**
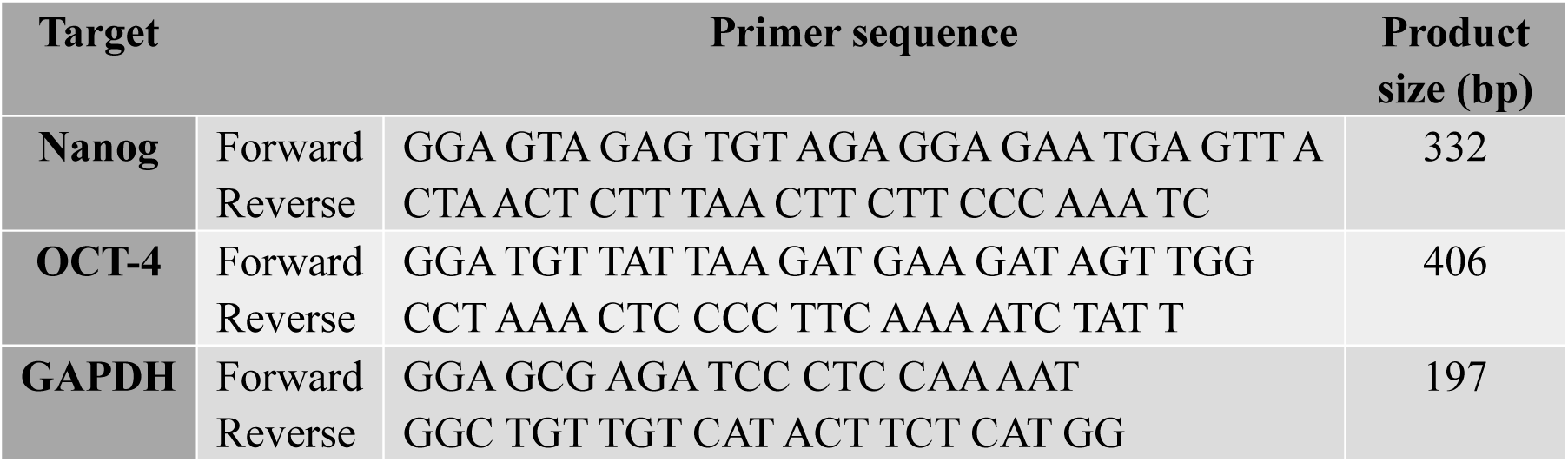
Primer sequences for pluripotency.

**Fig. 8.**
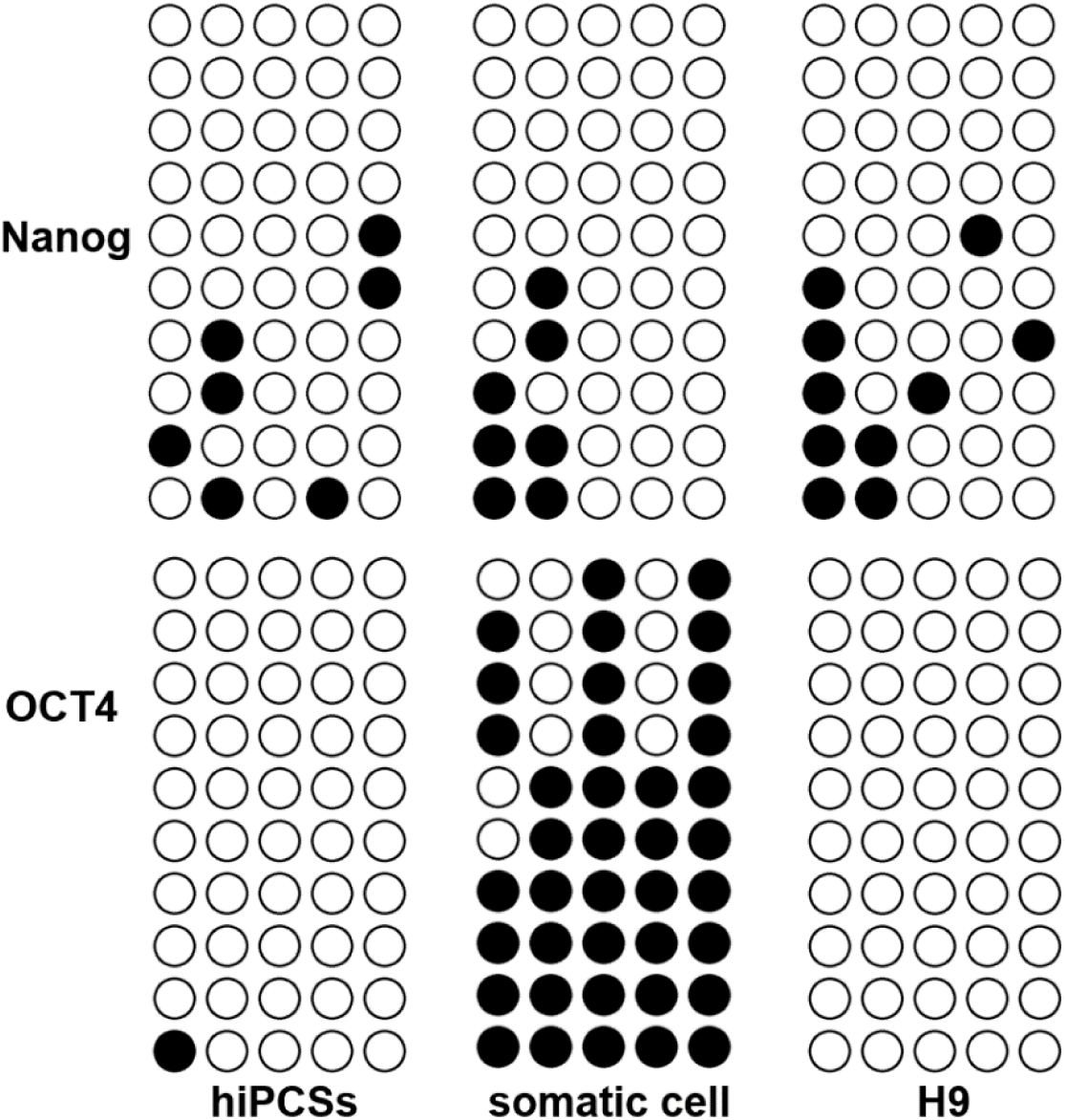
Bisulfite sequencing analysis of OCT4 and NANOG promoter regions in hiPSC, somatic cell and H9 (ESCs). Opened and closed circles indicate unmethylated and methylated CpGs, respectively.

### Directional differentiation of hiPSCs cell line

#### Culture hiPSC

The hiPSC was cultured in PSC easy human pluripotent stem cell (ESC/iPSC)culture medium (Cellapy:Cat. no. CA1001500/CA100110 0). The cells were subcultured for differentiated when the confluence reached 80%. We used Psceasy complete medium and Cell culture level 0.5mM EDTA subculture working fluid (Cellapy:Cat. no. CA3001500) to subculture hiPSCs.

#### Differentiation of iPSCs into neural stem cells (NSCs)

The cells were differentiated when the confluence reached 100%. PSC easy complete medium was removed and washed with 1ml PBS (Hyclone: Cat. no. SH30256.01). Added 3ml nerve stem differentiation complete medium to each pore cell, cultured in a 37°C constant temperature incubator with 5% CO_2_, after 48h added 2ml nerve stem differentiation complete medium again, continuous cultured for 48 hours, and then discarded 2ml supernatant, the complete culture medium of nerve stem differentiation of 2ml was added again, cultured in incubator for 48h. Microscopically, when Rosette structure appears (Fig.9A), cells can be digested for suspension culture. The cells were separated by Human Neural Stem Cells Digestible Fluid, then resuspended in Neural Trunk Inoculation Medium. The cells were cultured in Human Neural Stem Cell Maintenance Medium for 4-6 days and then subcultured (Fig.9B).

**Fig. 9.**
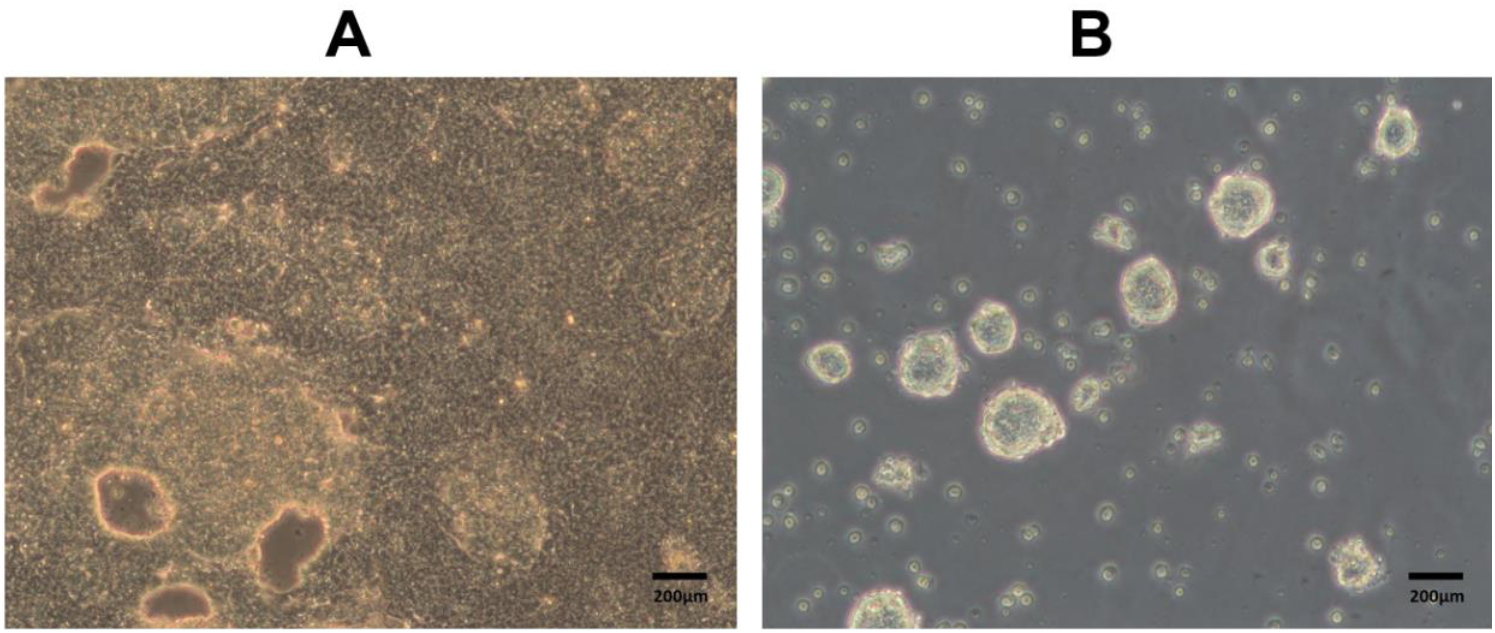
Differentiation of iPSCs into neural stem cells. (A) Rosette structure appeared. (B) Inoculation and maintenance culture of neural stem cells. Scale bar: 200 µm.

#### Differentiation of NSCs into neurons

Neural stem cells, which were subcultured 2-3 times, were collected in a 15 ml tube centrifuging 3 mins at 800rpm. Discarded the supernatant, added 1 ml Human Nerve Cell Digestive Juice, digested at 37°C for 15 mins. Discarded the supernatant after centrifuged at 1000×g for 5 mins, then resuspended in Human Nerve Differentiation Inoculation Medium. Next day, replaced the medium with the Human Nerve Differentiation and Maintenance Medium when the cells sticked to wall under microscope. Replaced the medium every two days, and neurofilaments could be observed after about 5-9 days (Fig.10A).

**Fig. 10.**
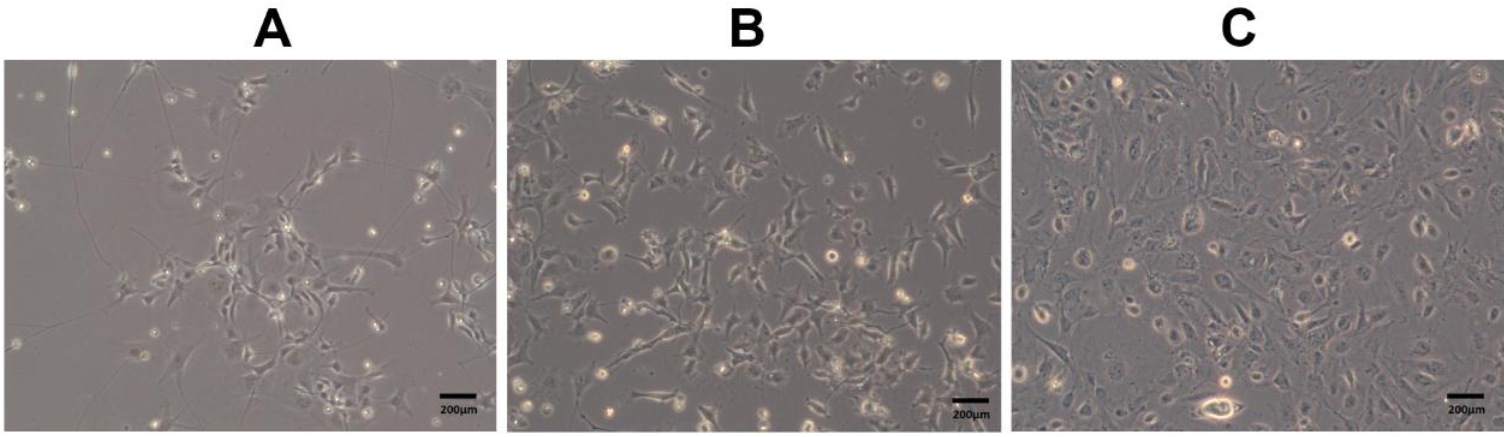
Differentiated cells. (A) Differentiation of neural stem cells into neurons. (B) Differentiation of neural stem cells into astrocyte. (C) Differentiation of hiPSCs into microvascular endothelial cells. Scale bar: 200 µm.

#### Differentiation of NSCs into astrocyte

Similar to neuronal cell differentiation, slightly different. After the neural stem cells were digested, the Human Neural Astrocyte Differentiation Medium resuspended the cells. Replaced the medium every two days, target cells were obtained in about seven days (Fig.10B).

#### Differentiation of hiPSCs into microvascular endothelial cells (MECs)

Digested hiPSCs with EDTA when the confluence reached 80%. Resuspended cells were counted and inoculated into 6-well plate according to 3 ×10^4^ cells/cm^2^. Added 10 uMY27632 and cultured with PSC easy complete medium. After 48h, absorbed the PSC easy complete medium and added the induction medium prepared in advance according to the 2 ml/ hole. After 5days, the medium was replaced by EC+RA medium. Microvascular endothelial cells were obtained after about 2days (Fig.10C).

### Identification of differentiated cells

Immunofluorescence staining were performed to identify the morphological feature of differentiated cells. Cells were fixed in 4% paraformaldehyde for 30 mins at 4°C. After washing with PBS, the cells were permeabilized with 0.3% Triton X-100 for 20 min, blocked in 1% bovine serum albumin for 40 mins and incubated with primary antibodies at room temperature. Cell nuclei were labeled with 4′,6-diamidino-2-phenylindole (DAPI, Vector, 20 mins, room temperature). For detection purposes, different fluorescently labelled secondary antibodies were selected depending on the specifications of the manufacturer. The antibodies were list in Table 1. Images were acquired using a fluorescence microscope (Carl ZeiSS, Germany) (Fig.11).

**Fig. 11.**
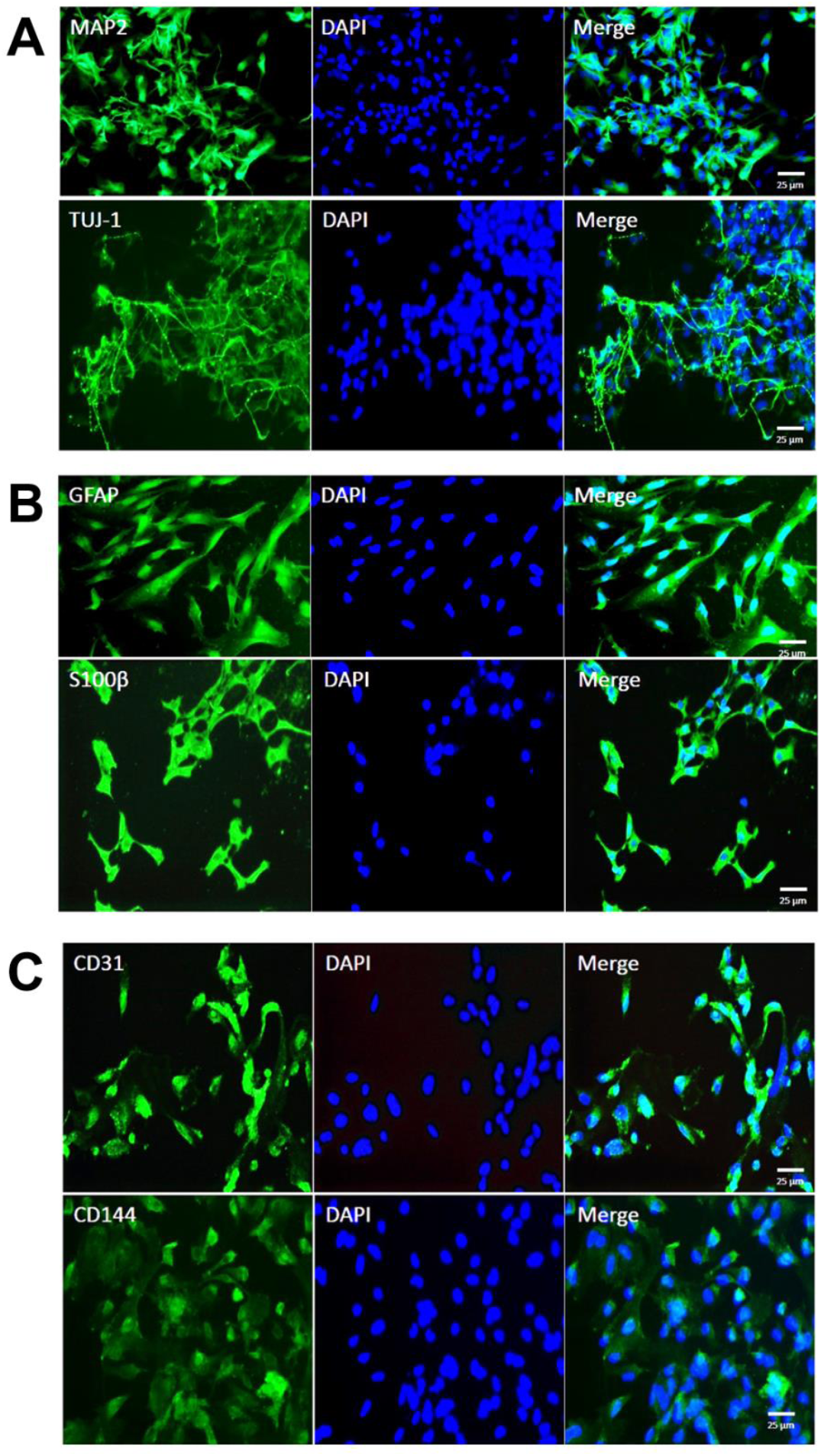
Immunofluorescence staining of differentiated cells. (A) Neurons were stained by MAP2 and TUJ-1. (B) Astrocytes were stained by GFAP and S100β. (C) Microvascular endothelial cells were stained by CD31 and CD144. Scale bars:25 µm.

## Results

### Characterization of hiPSCs and analysis of pluripotency

We utilized immunofluorescence and AP staining to evaluate the presence of specific pluripotency markers. Differentiating iPSC lines were observed the expression of human embryonic stem cell (hESC)-specific transcription factors NANOG, OCT4 and SOX2, as well as cell surface markers SSEA4, TRA-1-60 and TRA-1-81. hiPSC lines showed positive immunofluorescence labelling for above six hESC-specific antigens. (Fig. 3). If iPSC lines can maintain cellular morphological features and continued to express constant level of those markers after repeatedly passages, they were considered to be stable. Meanwhile, the pluripotency of hiPSC colonies displayed positive reaction for AP staining (Fig. 4). Through highly sensitive RT-PCR methods, the Sendai viral sequences were undetectable in reprogrammed iPSCs after 10 passages (Fig.5). All hiPSC lines were tested to be normal (46 chromosomes, XY) by GTG-banding karyotypic analysis (Fig.6). To further demonstrate the pluripotent status of hiPSCs in vivo, cells were subcutaneously injected CB-17 SCID male mice. All laboratory mice generated teratomas containing representative cell types of all three germ layers, as shown by H&E staining (Fig.7).

To identify the epigenetic modification in reprogrammed hiPSCs, we examined the methylation level of CpG dinucleotides in promoter regions of ESC-specific genes OCT4 and NANOG by bisulfite genomic sequencing (Fig.8). In T2DM patient, the CpG methylation ratios in OCT4 and NANOG promoters of hiPSCs were 2% and 14%, which almost corresponded to 0% and 12% in H9 (ESCs). Moreover, the CpG methylation ratio in OCT4 promoter regions of somatic cells was 78% in T2DM patient, respectively, significantly higher than that of hiPSCs and H9. Results revealed that the OCT4 and NANOG promoter regions were demethylated in hiPSCs relative to the somatic cells and were thus similar to those in hESCs. These results further indicated that the reprogrammed hiPSCs preserved ESC-like properties.

### Characterization of neuron, astrocyte and microvascular endothelial cell

NSCs derived from hiPSCs can proliferate and differentiate into both neural and glial lineage as stained by specific neuron markers MAP2, TUJ-1 (Fig.11A) and astrocyte markers GFAP, S100β (Fig.11B). Microvascular endothelial cells differentiated from hiPSCs were stained by CD31 and CD144 (Fig.11C). Immunocytochemical staining of above three cell types indicated cells stain positive for their specific markers, suggesting high reliability of our study.

## Discussion

Currently, a dominant challenge to our comprehension of the pathogenic mechanisms of diabetic brain injury has been the lack of physiologically relevant in vitro models which capture the precise patient genotype and phenotype, with the cell type of interest and specific protein expression. Induced pluripotent stem cells (iPSCs) technology, together with further cellular differentiation, supplies an attractive methodology for modeling neurological diseases, as well as for relevant drug discovery. With comparison to mouse counterparts, human iPSCs are considered to not only hold the similar self-renewal and pluripotency capacities of human embryonic stem cells (hESCs), but generate a limitless supply of targeted cell types for in vitro researches. Our previous studies and a series of clinical and basic researches on DM indicated that diabetic and hyperglycemia metabolic dysfunction leaded to various degrees of cognitive impairment and damage of learning and memory abilities(Wang et al., 2014, Lin et al., 2013, Jafari *et al*., 2009, Zhou *et al*., 2017, Lee *et al*., 2018). Substantial evidence have observed that there were dramatic pathological neuronal changes in streptozotocin (STZ) diabetes rats/mice, including neuronal apoptosis and synaptic alterations(Cai *et al*., 2018, Sadeghi *et al*., 2016), reduced expression levels of glial cell line-derived neurotrophic factor (GDNF) which was discovered as a survival neurotrophic factor to potently promotes the survival of many types of neurons(Sadeghi et al., 2016), increased expression of astrocyte glial fibrillary acidic protein (GFAP)(Sadeghi et al., 2016, Baydas *et al*., 2003). Moreover, diabetes and continuous high blood glucose could directly induce a vast spectrum of injuries of endothelial cells in cerebral vascular system(Xu *et al*., 2016), damage the permeability of BBB by down-regulating the expression of occluding and claudin-5(Yoo *et al*., 2016). Therefore, here we successfully created hiPSC-derived neuron (N), astrocyte (A) and microvascular endothelial cell (E), which displayed typical morphological characteristics and can be used to reveal the pathogenesis on molecular level and explore the innovative treatment strategies based on drug screening or other methods to target diabetic brain injury. In 2015, the Obama administration announced the plan for precision medicine initiative by the National Institutes of Health(Collins and Varmus, 2015, McCarthy, 2015). Precision medicine is defined as an approach to disease treatment and prevention targeted to maximize effectiveness on the basis of individual characteristics of genes, phenotype, environment, and lifestyle. As we know, DM results from a complex physiologic process that is determined by a large number of risk susceptibility genes and compounded environmental factors. Personalized model of brain injury with T2DM based on patient-specific iPSCs can avoid the genetic background issue and predict drug response of individual, which help advance the field of precision medicine. With the precision medicine initiative, it is clear that the generation of patient-derived iPSC technology will make it possible which can reserve a patient’s genetic and molecular background, and paved the way toward precision medicine goals of utilizing individual data to diagnose and therapy.

The next step of the study should unravel diabetic encephalopathy pathogenesis by utilizing these differential disease-relevant cell types (N, A and E). Co-culturing these three cells together in a 3D system, combined with advanced microfluidic chip technology, to further create a brand new personalized human brain cell models in vitro, which will provide a new way to accurately simulate the pathogenic microenvironment of diabetic brain injury, study the identification, intervention and evaluation of cellular therapeutic targets, and further investigate the disease-specific mechanisms to achieve further insights into disease pathogenesis and establish individualized-targeted therapy.

## Footnotes

### Author contributions

Conceptualization and Methodology: W.L., Y.-Z.L.; Software, Validation, Formal analysis, Investigation, Data curation and Visualization: W.L.; Writing: W.L., J.T. Resources: P.Z.; Supervision, Project administration and Funding acquisition: Y.-Z.L.

